## Extended Data for "Mechanism of NanR gene repression and allosteric induction of bacterial sialic acid metabolism"

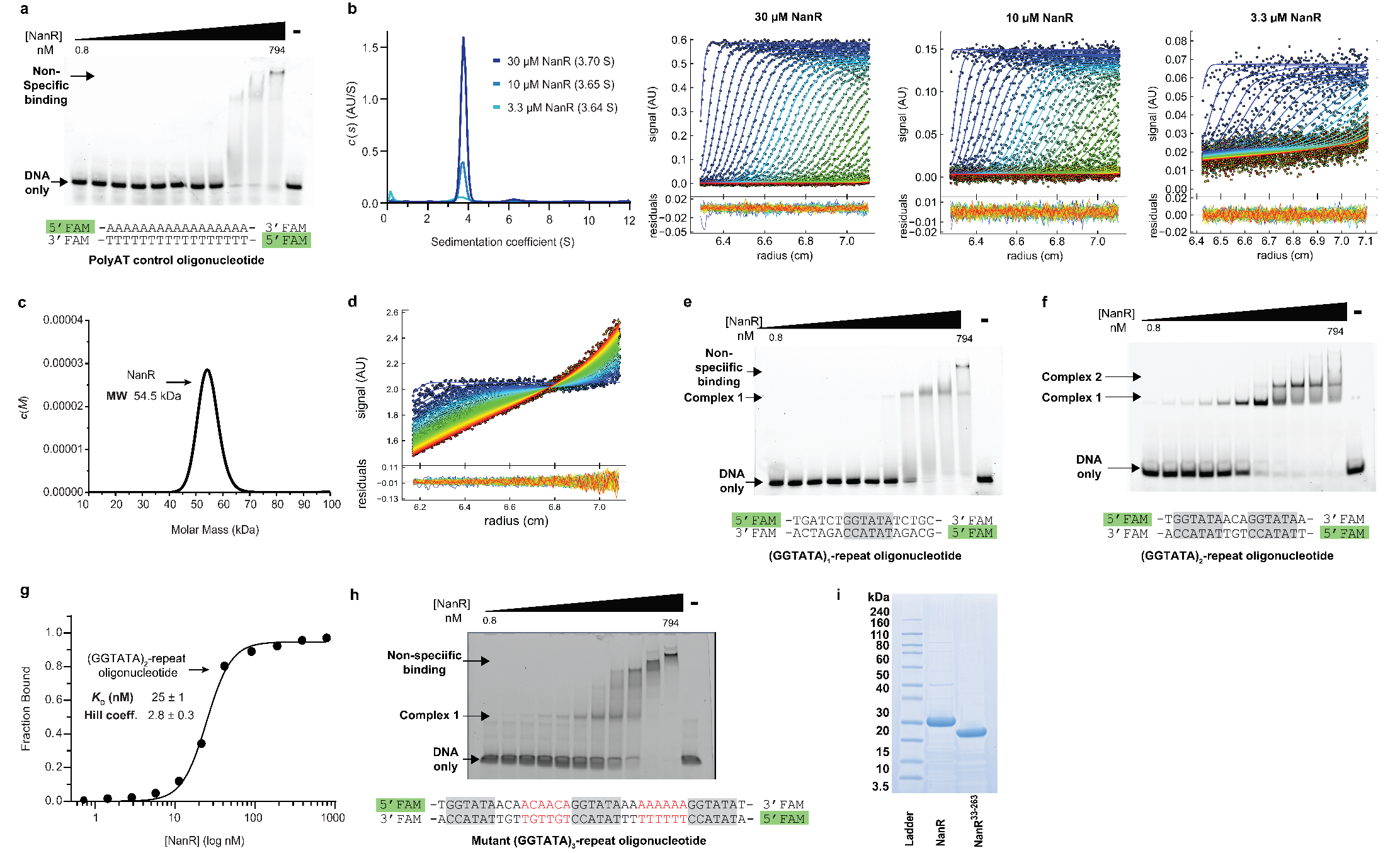

***Extended Data* Fig 1 | a,** Specific DNA binding is abolished in the poly adenine/thymine (poly-AT) control oligonucleotide. At high concentrations (>200 nM), non-specific binding is observed. **b,** Sedimentation velocity data for NanR at 3.3-30 µM is a single monodisperse species at 3.64-3.70 S. The raw data and residual fits are shown to the right of the pane, where every third scan is shown. The data are fit to a continuous size [*c*(*s*)] distribution, as implemented in SEDFIT^1^. The hydrodynamic properties for the fit are listed in ***Extended Data* Table 1. c,** Sedimentation velocity data for NanR at 10 µM fit to a continuous mass [*c*(M)] distribution giving a molar mass of 54.5-kDa across the peak, consistent with the calculated mass of the NanR dimer (59.0-kDa). The r.m.s.d. for the fit was 0.004. **d,** When FAM alone is mixed with NanR at 10 µM and sedimentation velocity data is collected at 50,000 rpm and 495 nm (monitoring only FAM), the FAM molecule is not transported with the protein. This control experiment demonstrates that NanR does not bind the FAM label. **e,** Poor DNA binding activity is observed with a single GGTATA repeat, suggesting the regulation mechanism is dependent on cooperative binding. **f,** In contrast, two concentration-dependent complexes are observed when two GGTATA repeats are present. The concentration of DNA in the experiments **e** and **f** is 10 nM. **g,** The binding isotherm from the (GGTATA)_2_-repeat oligonucleotide with NanR. The data was best fit to a Hill equation (AIC value of 99%), when compared with the Mass Action Law (AIC value of 1%). **h,** Increasing the length of the spacer between the GGTATA repeats altered NanR binding. The additional nucleotides are highlighted red in the mutant (GGTATA)_3_-repeat oligonucleotide. The concentration of DNA in the experiment is 10 nM. **i,** SDS-PAGE analysis showing protein molecular weight ladder (Lane 1), and purified *E. coli* NanR (Lane 2) and NanR^33–263^ (Lane 3) following size exclusion chromatography.

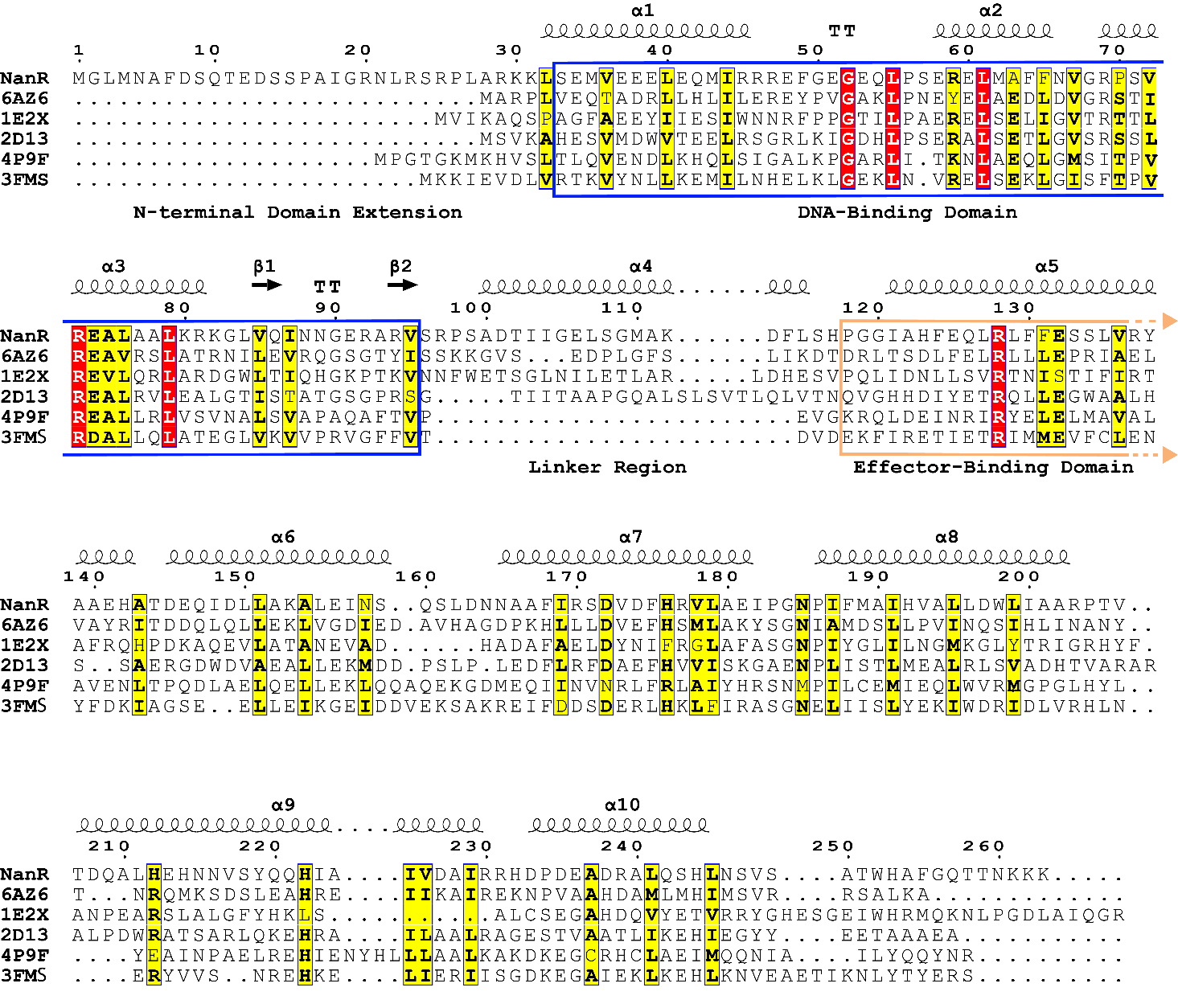
***Extended Data* Fig. 2 | NanR has a longer N-terminal sequence, when compared to known GntR proteins.** Multiple sequence alignment performed using Clustal Omega^2^ and generated using ESPript 3.0^3^. The top five hits from a sequence homology search within the PDB were used in the sequence alignment^4-8^. The red background highlights identical residues, while the yellow background indicates similar residues. Secondary structure elements depicted above the alignment were generated from the crystal structure of *E. coli* NanR (**Fig. 4**), solved in this study. The N-terminal DNA-binding domain (blue box), the linker region, and the C-terminal effector-binding domain (beige box) are highlighted.

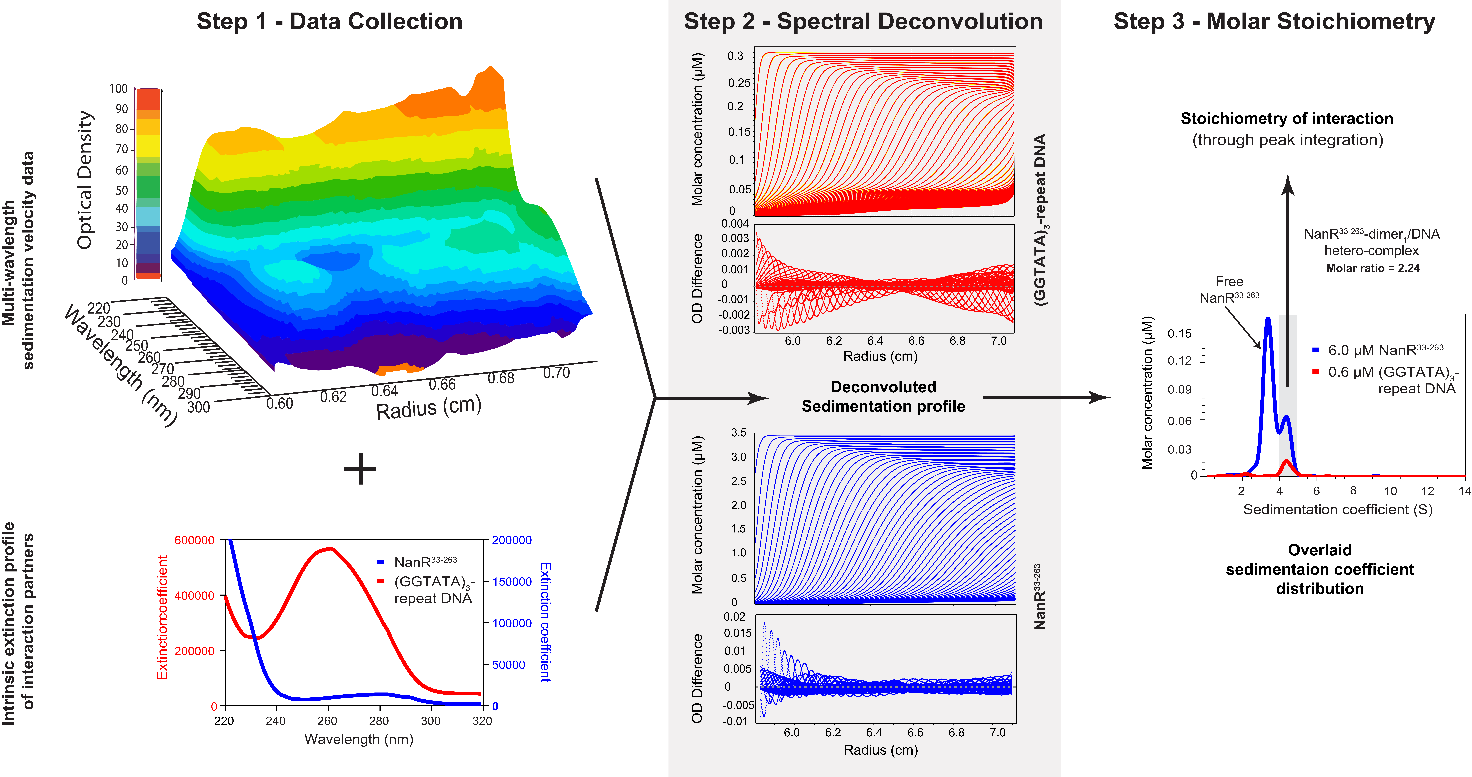
***Extended Data* Fig. 3 | Multiwavelength sedimentation velocity data for NanR^33^**^–^**^263^ and DNA and analysis workflow. *Step 1,*** four-dimensional multiwavelength sedimentation velocity data (top left) is collected across a range of wavelengths (220–300 nm). Only one time point is presented, highlighting peaks at 220 nm and 260–280 nm, consistent with a mixture of protein and DNA. An intrinsic extinction profile (bottom left) is generated for each interaction partner by globally fitting a dilution series of absorbance profiles using UltraScan^9^. ***Step 2****,* the intrinsic extinction profile is then used to spectrally deconvolute the multiwavelength data into separate sedimentation profiles for each interacting component (NanR^33–263^ in red, (GGTATA)_3_-repeat DNA in blue) by employing the non-negatively constrained least squares algorithm^9,10^. This process also scales the data to molar concentrations. ***Step 3,*** once deconvoluted and on a molar scale, the stoichiometry of the complex can simply be extracted by integrating the molar ratio of the co-migrating peaks (shaded box). In addition to this spectral characterization, the hydrodynamic information can also be accessed, providing such parameters as the molar mass and frictional ratio of each species. Together this enables interacting species to be characterized with high resolution.

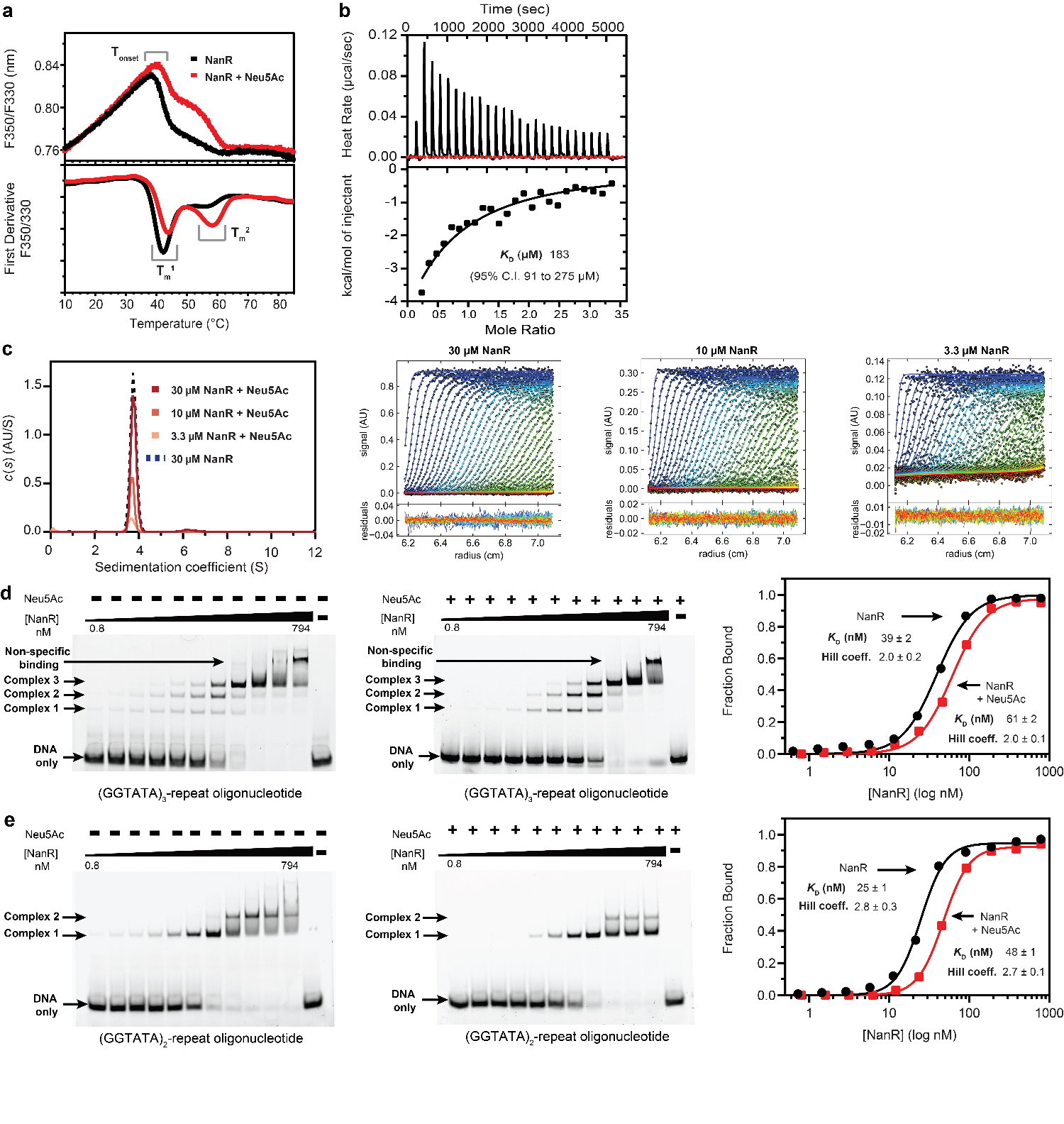
***Extended Data* Fig. 4 | Binding interaction between NanR and Neu5Ac and effect on DNA binding. a,** Thermal stability of NanR (33 µM in black) with and without Neu5Ac (20 mM in red) using differential scanning fluorimetry. The results are presented as the ratio of intrinsic tryptophan fluorescence at the emission wavelengths of 350 and 330 nm (top pane) and the first derivative of this fluorescence ratio (lower pane). An increase in the thermal stability of NanR is observed in the presence of Neu5Ac at both the T_onset_ and the first transition melting temperature (T_m_^1^), which verifies Neu5Ac can bind NanR. The second transition melting temperature (T_m_^2^), observed only in the presence of Neu5Ac, may reflect increased thermal stability of the C-terminal effector-binding domain of NanR when it binds Neu5Ac. **b,** ITC isotherm of Neu5Ac (2 mM) titrated into NanR (120 μM) in buffer C at 25 °C. The titration involved 25 injections of ligand solution (2 μL) into the protein sample cell with a 200 sec interval between subsequent injections. When fitted, the data gave a *K*_D_ value of 183 µM. **c,** Sedimentation velocity data for NanR in the presence of Neu5Ac (20 mM) at 3.3–30 µM is a single monodisperse species at 3.64–3.70 S. The raw data and residual fits are shown to the right of the pane, where every third scan is shown. The data is fit to a continuous size [*c*(*s*)] distribution, as implemented in SEDFIT^1^. The hydrodynamic properties for the fit are listed in ***Extended Data* Table 1. d,** Three concentration-dependent complexes (1–3) are observed by EMSA when NanR (0.8–794 nm) is titrated against FAM-5’-labeled DNA (10 nM) with three GGTATA binding sites (left pane). In the presence of Neu5Ac (20 mM), a small decrease in DNA binding affinity is observed (center pane). Binding isotherm from the EMSA experiment (right pane) for NanR only (black) and in the presence of Neu5Ac (red). The data was best fit to the Hill equation. **e,** Two concentration-dependent complexes (C1-2) are observed by EMSA when NanR (0.8–794 nm) is titrated against FAM-5ʹ-labeled DNA (10 nM) with two GGTATA binding sites (left pane). In the presence of Neu5Ac (20 mM), a small decrease in DNA binding affinity is observed (center pane). Binding isotherm from the EMSA experiment (right pane) for NanR only (black) and in the presence of Neu5Ac (red). The data was best fit to the Hill equation.

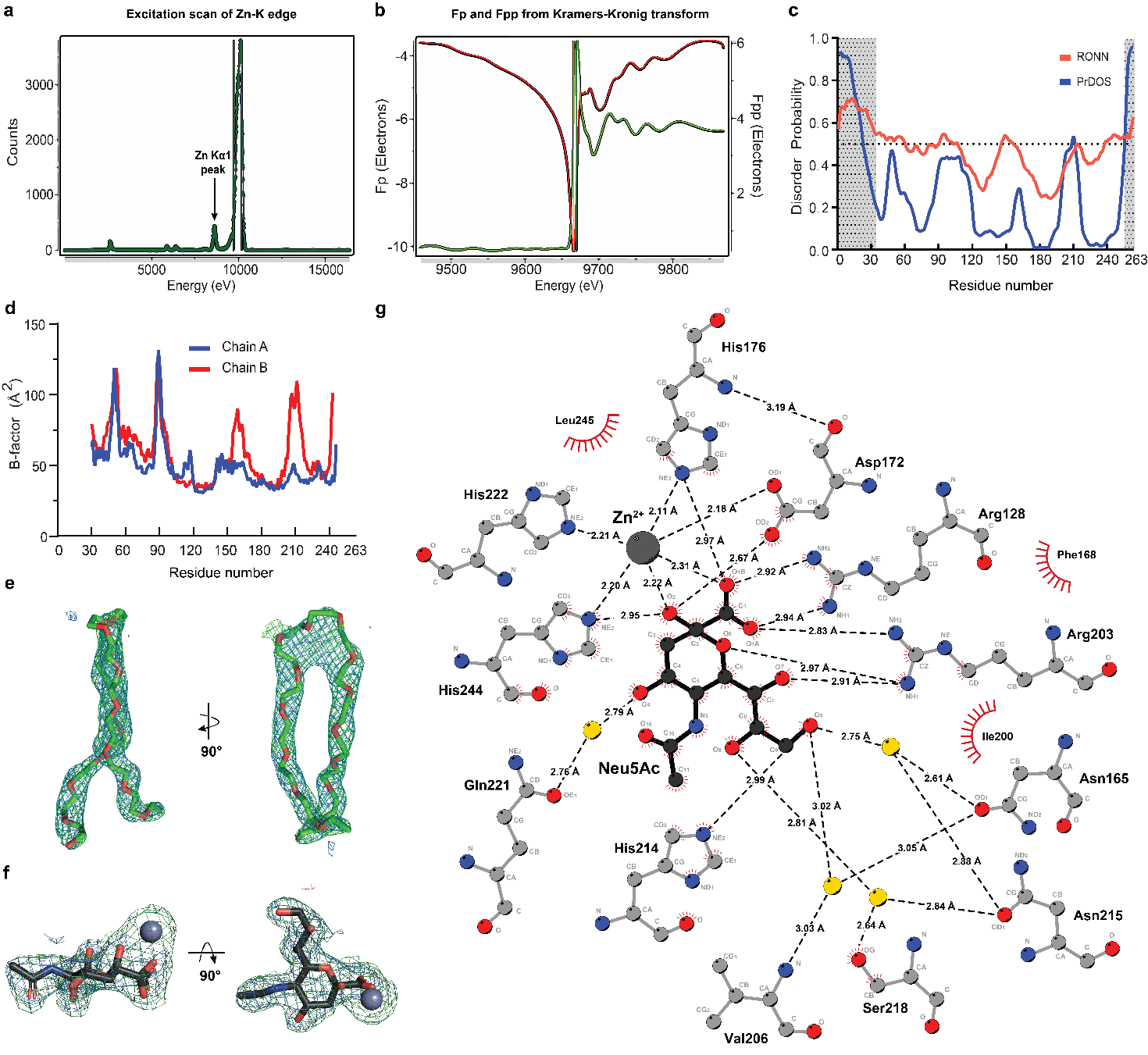
***Extended Data* Fig. 5 | Detection of elemental zinc, disorder prediction, and ligand analysis of the crystal structure. a,** Elemental analysis of the NanR crystal using X-ray fluorescence suggests the presence of zinc (Zn Kα_1_ peak corresponding to an emission energy of ~8.6 KeV^11^). **b,** MAD scan of the NanR crystal. Derivation of f’ (red) and f” (green) from the normalized MAD scan was calculated using the Kramers-Kronig transform in Chooch^12^ The absorption edge of Zn corresponds to 9670.10 eV and the inflection energy to 9665.92 eV. **c,** Disorder probability for NanR estimated from the RONN (https://www.strubi.ox.ac.uk/RONN) and PrDOS (http://prdos.hgc.jp/cgi-bin/top.cgi) online servers. The regions of disorder are highlighted (grey with dot). **d.** Plot of B-factor values for chains A (Neu5Ac/Zn^2+^ bound, blue) and B (ligand free, red) of the NanR crystal structure. **e,** Omit map showing the electron density for polyethylene glycol. Here, the 2F*o*-F*c* electron density map (1.0 σ, blue mesh) and the mF*o*-F*c* omit electron density map (3.0 σ, green mesh) is shown. **f,** Omit map showing the electron density for Neu5Ac in its β-anomeric form (depicted as sticks) and Zn^2+^ (grey sphere). Here, the 2F*o*-F*c* electron density map (1.0 σ, blue mesh) and the mF*o*-F*c* omit electron density map (3.0 σ, green mesh) is shown. **g,** Schematic representation of the NanR-Neu5Ac-Zn^2+^ interaction network. The hydrogen-bonded network is presented (black dashed lines) and the distances between partners are labeled. Key interacting water molecules (yellow spheres) and hydrophobic contacts (red arcs with spokes) are shown. Figure generated by LigPlot+^13^.

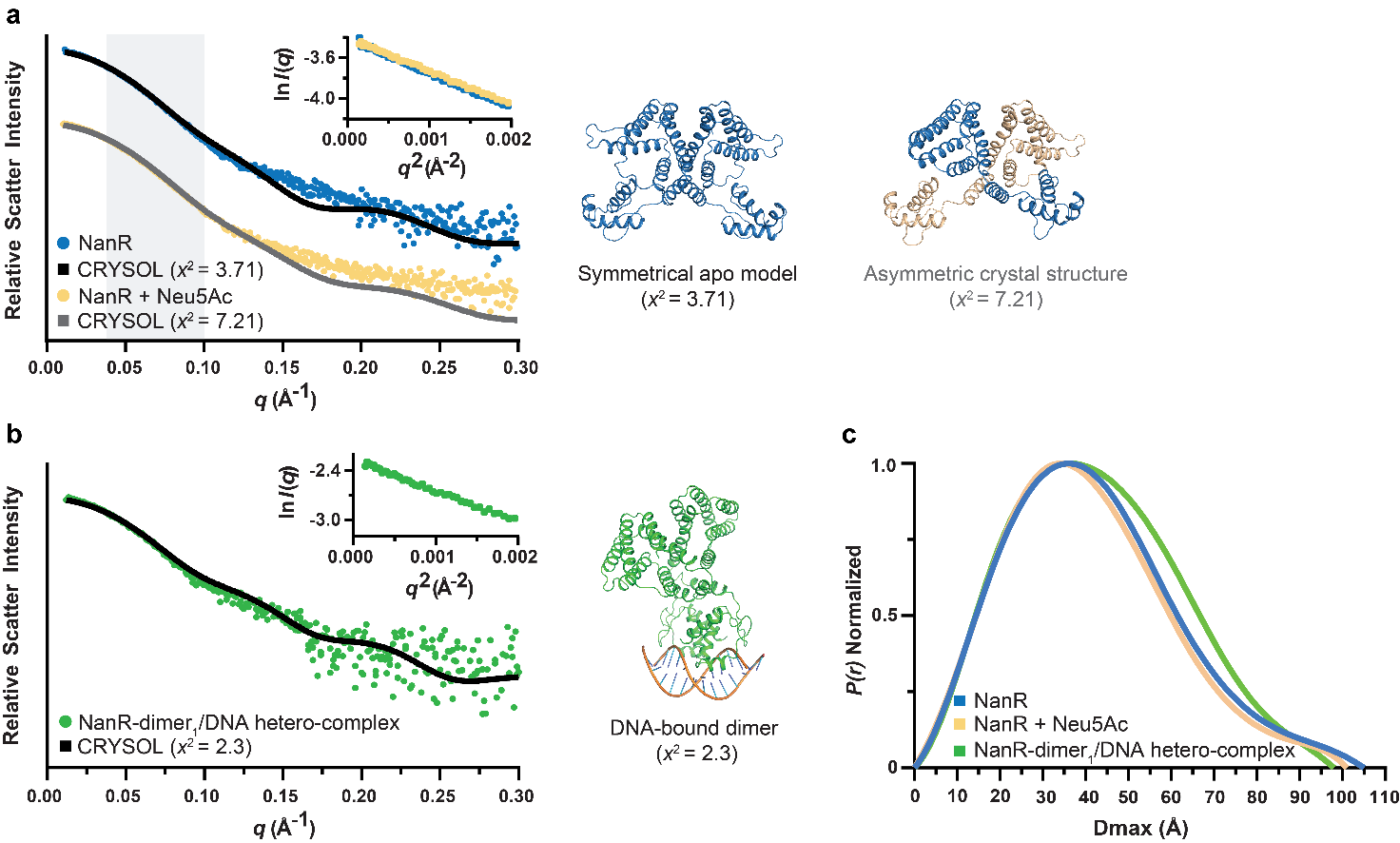
***Extended Data* Fig. 6 | SAXS for *E. coli* NanR and the NanR-DNA hetero-complex. a,** Small angle X-ray scattering data for NanR alone (blue) and NanR in the presence of Neu5Ac (beige). The difference at low *q* is highlighted in the shaded area. The scattering data for NanR (blue) is best fit (black line, Χ^2^=3.71) to the back-calculated scattering pattern of the symmetrical Neu5Ac-free dimer (shown in cartoon, blue), suggesting that the Neu5Ac-free structure is a reasonable model for NanR in solution. The scattering data for NanR in the presence of Neu5Ac (beige) is best fit (grey line, Χ^2^=7.21) to the back-calculated scattering pattern for the asymmetric Neu5Ac-bound dimer (shown in cartoon, blue and beige), supporting a conformational change upon Neu5Ac binding. **b,** Small angle X-ray scattering data for the NanR-dimer_1_/DNA hetero-complex (green). The back-calculated scattering pattern based on the cryo-EM NanR-dimer_1_/DNA hetero-complex structure (shown in cartoon, green) fits well to the experimental data (black, Χ^2^ =2.3), suggesting that the cryo-EM model is analogous to the solution structure. **c,** The pairwise distribution plots, calculated from the scattering data, estimate the maximum inter-particle dimension (D_max_) for NanR (blue), NanR in the presence of Neu5Ac (beige) and the NanR-dimer_1_/DNA hetero-complex (green). For all data, the Guinier plots, inset (respective colors), are linear, showing that the sample is free from aggregation or inter-particle interference.

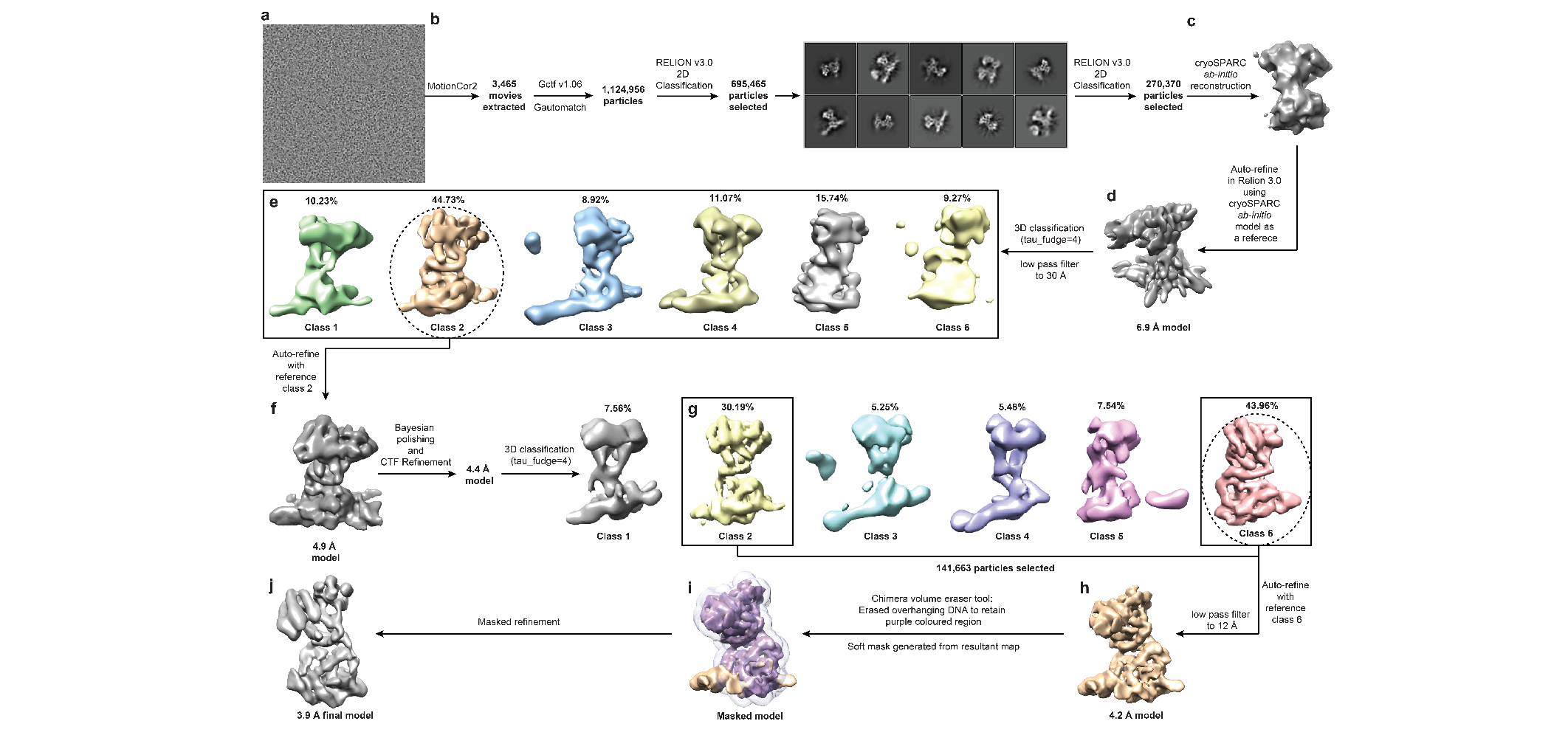
***Extended Data* Fig. 7 | Cryo-EM processing pipeline for the NanR-dimer_1_/DNA hetero-complex. a,** Electron micrograph at 2.4 µm defocus. **b,** Two rounds of 2D classification retaining 270,370 particles that had high signal-to-noise. **c,** Initial 3D *ab initio* reconstruction. **d,** The first round of auto-refinement resulted in a 6.9 Å reconstruction. **e,** Initial 3D classification with a low pass filter to 30 Å generated a total of six discrete classes, where four of these showed features consistent with DNA binding (black box). Class 2 has the highest signal-to-noise (dash circle). The abundance of each class is shown by percentage. **f,** The second round of auto-refinement, using class 2 as a reference resulted in a 4.9 Å reconstruction. **g,** Following Bayesian polishing and CTF refinement, a second 3D classification generated a total of six discrete classes. Classes 2 and 6 have the highest signal-to-noise (black box). The abundance of each class is shown by percentage. **h,** The third round of auto-refinement, using class 6 as a reference (dash circle in **g**) and a low pass filter to 12 Å resulted in a 4.2 Å reconstruction. **i,** A soft mask was generated around the core of the model to exclude the regions with the highest flexibility (grey density). **j,** The masked refinement resulted in a final 3.9 Å reconstruction of the NanR-dimer_1_/DNA hetero-complex.

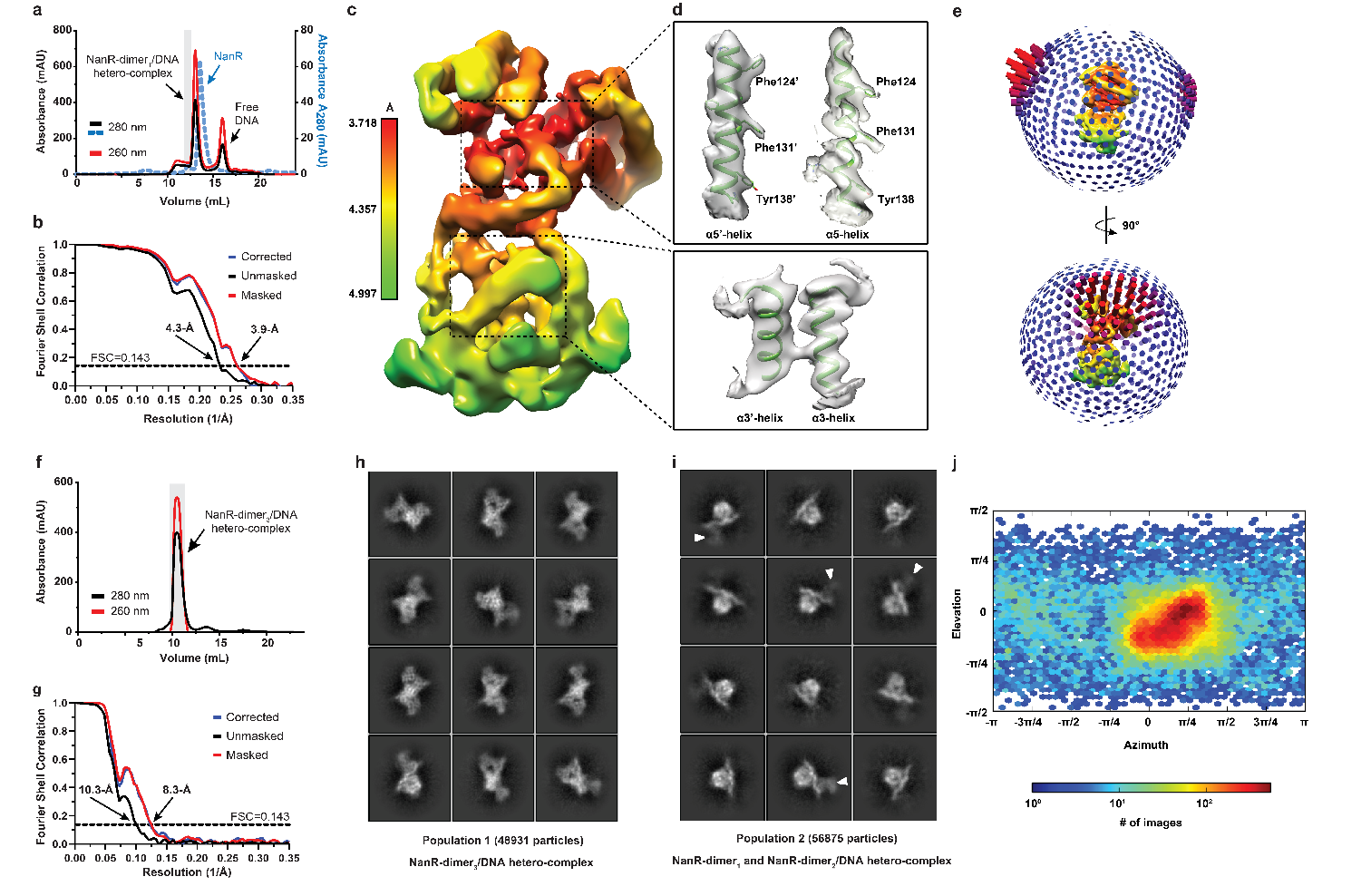
***Extended Data* Fig. 8 | Sample preparation, map quality, and preferred orientation of the NanR-dimer_1_ and NanR-dimer_3_/DNA hetero-complex** **cryo-EM structures. a,** Size exclusion chromatogram highlighting the purification of the NanR-dimer_1_/DNA hetero-complex from free NanR and DNA. The shaded area was pooled and subsequently used for cryo-EM sample preparation. **b,** Fourier shell correlation plot indicating a resolution of 3.9 Å and 4.3 Å for the masked (red) and unmasked (black) 3D reconstructions of the NanR-dimer_1_/DNA hetero-complex, respectively, as reported by the FSC=0.143 criterion. **c,** Local resolution estimation for the final masked 3D reconstruction of the NanR-dimer_1_/DNA hetero-complex. The color bar outlines the resolution range of the map in Å. **d,** Helices α5, which contribute to the dimer interface (top inset), and helices α3, which bind DNA at the major groove (lower inset), present the highest level of resolvability. Sidechains for residues that could be assigned are labeled and shown as sticks. **e,** Angular distribution estimation for the NanR-dimer_1_/DNA hetero-complex dataset suggests a preferential orientation of the particles. Each cylinder shown in the distribution represents one view, where the height is reflective of the number of particles in that view. **f,** Size exclusion chromatogram highlighting the purification of the NanR-dimer_3_/DNA hetero-complex from free NanR and DNA. The shaded area was pooled and subsequently used for cryo-EM sample preparation. **g,** Fourier shell correlation plot indicating a resolution of 8.3 Å and 10.3 Å for the masked (red) and unmasked (black) 3D reconstructions of the NanR-dimer_3_/DNA hetero-complex, respectively, as reported by the FSC=0.143 criterion. **h,** 2D class averages from population 1 comprised only NanR-dimer_3_/DNA hetero-complex particle projections. **i,** 2D class averages from population 2 are consistent with a mixture of NanR-dimer_1_ and NanR-dimer_2_/DNA hetero-complex particle projections. The second NanR dimer is highlighted by a white arrow. **j,** Angular distribution of NanR-dimer_3_/DNA hetero-complex particle projections suggests a preferential orientation according to cryoSPARC v2 non-uniform refinement^14^.

***Extended Data* Table 1 | Summary of sedimentation velocity analysis of NanR, NanR in the presence of Neu5Ac and DNA.**

|  | **Wavelength**  **(nm)** | | **Sedimentation coefficient**  **(S)** | **Molar mass**  **(kDa)** | **Calculated molar mass (kDa)** | **Frictional ratio**  **(*f*/*f*_0_)** | **Fit RMSD** |
| --- | --- | --- | --- | --- | --- | --- | --- |
| (GGTATA)_3_-repeat DNA (80 nM) | 495 (fluorescence) | | 3.00 | 23.0 | 22.6 | 1.68 | 11.3 |
| NanR (30 µM) | 280 | | 3.70 | 54.5 | 59.0 | 1.32 | 0.005 |
| NanR (10 µM) | 280 | | 3.65 | 54.2 | 59.0 | 1.32 | 0.004 |
| NanR (3.3 µM) | 280 | | 3.64 | 54.3 | 59.0 | 1.32 | 0.004 |
| NanR (10 µM) + FAM (3 µM) | 495 (absorbance) | | - | - | - | - | 0.004 |
| NanR (30 µM) + Neu5Ac (20 mM) | 280 | | 3.71 | 54.3 | 59.0 | 1.29 | 0.005 |
| NanR (10 µM) + Neu5Ac (20 mM) | 280 | | 3.67 | 54.3 | 59.0 | 1.31 | 0.004 |
| NanR (3.3 µM) + Neu5Ac (20 mM) | 280 | | 3.65 | 54.4 | 59.0 | 1.30 | 0.004 |
| **Fixed parameters** | |  | |  |  |  |  |
| Buffer density (g/cm^3^) | | 1.006 | | | | | |
| Buffer viscosity (cP) | | 1.027 | | | | | |
| Partial specific volume (NanR) | | 0.7295 | | | | | |
| Partial specific volume ((GGTATA)_3_-repeat DNA) | | 0.5500 | | | | | |

***Extended Data* Table 2 | Summary of fluorescence-detection sedimentation velocity analysis of NanR titrated against DNA.**

| **Sample** | **Weight-averaged sedimentation coefficient**  **(S) [Peak integration range, S]** | **Frictional ratio**  **(*f*/*f*_0_)** | **Fit RMSD** |
| --- | --- | --- | --- |
| (GGTATA)_3_-repeat DNA (80 nM) | 3.00 [2.0–3.5] | 1.68 | 11.3 |
| 0.8 nM NanR (+ 80 nM (GGTATA)_3_-repeat DNA) | 3.03 [3.0–12.0] | 1.70 | 9.4 |
| 1.6 nM NanR (+ 80 nM (GGTATA)_3_-repeat DNA) | 3.04 [3.0–12.0] | 1.71 | 10.0 |
| 3.1 nM NanR (+ 80 nM (GGTATA)_3_-repeat DNA) | 3.04 [3.0–12.0] | 1.72 | 8.9 |
| 6.2 nM NanR (+ 80 nM (GGTATA)_3_-repeat DNA) | 3.07 [3.0–12.0] | 1.72 | 10.1 |
| 12.4 nM NanR (+ 80 nM (GGTATA)_3_-repeat DNA) | 3.14 [3.0–12.0] | 1.73 | 8.6 |
| 24.8 nM NanR (+ 80 nM (GGTATA)_3_-repeat DNA) | 3.47 [3.0–12.0] | 1.86 | 10.0 |
| 49.6 nM NanR (+ 80 nM (GGTATA)_3_-repeat DNA) | 4.09 [3.0–12.0] | 1.90 | 8.7 |
| 99.3 nM NanR (+ 80 nM (GGTATA)_3_-repeat DNA) | 6.00 [3.0–12.0] | 1.97 | 10.8 |
| 198.5 nM NanR (+ 80 nM (GGTATA)_3_-repeat DNA) | 8.28 [3.0–12.0] | 2.39 | 10.8 |
| 397.0 nM NanR (+ 80 nM (GGTATA)_3_-repeat DNA) | 8.50 [3.0–12.0] | 2.43 | 17.0 |
| **Fixed parameters** |  |  |  |
| Buffer density (g/cm^3^) | 1.006 | | |
| Buffer viscosity (cP) | 1.027 | | |
| Partial specific volume (NanR) | 0.7295 | | |
| Partial specific volume ((GGTATA)_3_-repeat DNA) | 0.5500 | | |

***Extended Data* Table 3 | Summary of sedimentation velocity analysis of NanR and NanR^33-263^ in the presence of DNA.**

| **Sample** | **Wavelength**  **(nm)** | **Weight-averaged**  **sedimentation coefficient (x10^-13^ S)** | **Frictional ratio**  **(*f*/*f*_0_)** | **Fit RMSD** |
| --- | --- | --- | --- | --- |
| (GGTATA)_3_-repeat DNA (3 µM) | 495 | 2.95 | 1.68 | 0.003 |
| 3 µM NanR (+ 3µM (GGTATA)_3_-repeat DNA) | 495 | 3.80 | 1.51 | 0.003 |
| 12 µM NanR (+ 3µM (GGTATA)_3_-repeat DNA) | 495 | 6.32 | 1.56 | 0.003 |
| 24 µM NanR (+ 3µM (GGTATA)_3_-repeat DNA) | 495 | 9.11 | 1.62 | 0.003 |
| 10 µM NanR^33-263^ | 495 | 3.38 | 1.25 | 0.004 |
| 3 µM NanR^33-263^ (+ 3µM (GGTATA)_3_-repeat DNA) | 495 | 3.43 | 1.61 | 0.003 |
| 12 µM NanR^33-263^ (+ 3µM (GGTATA)_3_-repeat DNA) | 495 | 4.39 | 1.53 | 0.003 |
| 24 µM NanR^33-263^ (+ 3µM (GGTATA)_3_-repeat DNA) | 495 | 4.45 | 1.49 | 0.003 |
| **Fixed parameters** |  |  |  |  |
| Buffer density (g/cm^3^) |  | 1.006 | | |
| Buffer viscosity (cP) |  | 1.027 | | |
| Partial specific volume (NanR) |  | 0.7295 | | |
| Partial specific volume (NanR^33–263^) |  | 0.7300 | | |
| Partial specific volume ((GGTATA)_3_-repeat DNA) |  | 0.5500 | | |

***Extended Data* Table 4 | Integration results from multi-wavelength sedimentation velocity analysis of deconvoluted NanR^33-263^ and (GGTATA)_3_-repeat DNA sedimentation profiles.**

|  | **NanR^33-263^**  **only** | **(GGTATA)_3_-repeat DNA only** | **NanR^33-263^:DNA loading ratio** | |
| --- | --- | --- | --- | --- |
|  |  |  | **3:1** | **10:1** |
| Sed. Coefficient (×10^-13^ S) * | 3.42  (3.09, 3.77) | 2.76  (2.63, 2.89) | 4.39  (4.10, 4.67) | 4.46  (3.58, 5.33) |
| Dif. Coefficient (×10^-7^ D) * | 6.79  (4.09, 9.49) | 6.99  (6.10, 7.86) | 4.97  (4.03, 5.91) | 5.04  (3.39, 6.70) |
| Measured molar ratio ^‡^ | n/a | n/a | 2.44 | 2.24 |
| Weight-averaged $\bar{v}$ (mL g^-1^) ^§^ | 0.730 | 0.55 | 0.677 | 0.677 |
| Measured molar mass (kDa) ^¶^ | 45.3  (29.7, 62.8) | 21.4  (17.9, 24.8) | 66.3 | 66.4 |
| Theoretical molar mass (kDa) ^#^ | 51.3 | 21.5 | 72.8 | 72.8 |
| Oligomeric state of hetero-complex | n/a | n/a | NanR-dimer_1_/DNA | NanR-dimer_1_/DNA |
| * Sedimentation and diffusion coefficients observed following 2DSA-Monte Carlo analysis. Parameters are obtained from integration of pseudo-3D plots using UltraScan^15^. All measured values represent the mean from the Monte Carlo analysis. The values in parentheses are the 95% confidence intervals from the Monte Carlo analysis. | | | | |
| ^‡^ Partial concentration is determined from peak integration of the co-migrating species in both the NanR and DNA datasets. Because the data is scaled to molar concentrations, the molar ratio and thus stoichiometry of the hetero-complex can be inferred. | | | | |
| ^§^ Partial specific volume ($\bar{v}$) of the hetero-complex, estimated from the weight-average of the protein and DNA components. A $\bar{v}$ of 0.7295 mL g^-1^ was used for NanR, while a $\bar{v}$ of 0.55 mL g^-1^ was used for DNA. The equation used to calculate the weight-averaged $\bar{v}$ is presented in the Methods. | | | | |
| ^¶^ The measured molar mass is estimated based on the hydrodynamic parameters (sedimentation and diffusion coefficient) and the weighted-averaged $\bar{v}$, calculated using the measured molar ratio. The values in parentheses are the 95% confidence intervals from the Monte Carlo analysis for the pure components. | | | | |
| ^#^ The theoretical mass is predicted based upon the amino acid or nucleic acid sequence within UltraScan^15^. The mass of each hetero-complex is predicted based on the observed molar ratio. | | | | |

***Extended Data* Table 5 | Integration results from multi-wavelength sedimentation velocity analysis of deconvoluted NanR and (GGTATA)_3_-repeat DNA sedimentation profiles.**

|  | **NanR**  **only** | **(GGTATA)_3_-repeat DNA only** | **NanR:DNA loading ratio** | | | | | | |
| --- | --- | --- | --- | --- | --- | --- | --- | --- | --- |
|  |  |  | **1:1**  (Species 1) | **1:1**  (Species 2) | **3:1**  (Species 1) | **3:1**  (Species 2) | **6:1**  (Species 2) | **6:1**  (Species 3) | **10:1**  (Species 3) |
| Sed. Coefficient  (×10^-13^ S) * | 3.72  (3.50, 3.94) | 2.76  (2.63, 2.89) | 5.67  (5.09 6.26) | 7.45  (7.01, 7.90) | 5.62  (4.83, 6.41) | 7.46  (6.82, 8.11) | 7.54  (7.13, 7.95) | 7.99  (7.35, 8.61) | 8.31  (7.95, 8.68) |
| Dif. Coefficient  (×10^-7^ D) * | 6.3  (3.84, 8.76) | 6.99  (6.10, 7.86) | 4.37  (2.46, 6.30) | N/D† | 4.40  (2.18, 6.11) | N/D ^†^ | N/D ^†^ | 3.16  (2.65, 3.66) | 3.40  (2.49, 4.31) |
| Measured molar ratio ^‡^ | n/a | n/a | 2.45 | 4.25 | 2.37 | 3.63 | 4.65 | 6.44 | 6.21 |
| Weight-averaged $\bar{v}$  (mL g^-1^) ^§^ | 0.730 | 0.550 | 0.682 | 0.702 | 0.682 | 0.702 | 0.702 | 0.710 | 0.710 |
| Measured molar mass (kDa) ^¶^ | 54.0  (30.0, 77.9) | 21.4  (17.9, 24.8) | 98.9 | N/D† | 97.6 | N/D† | N/D† | 211.8 | 204.8 |
| Theoretical molar mass (kDa) ^#^ | 59.0 | 21.5 | 80.5 | 139.5 | 80.5 | 139.5 | 139.5 | 198.5 | 198.5 |
| Oligomeric state of  hetero-complex | n/a | n/a | NanR-dimer_1_/DNA | NanR-dimer_2_/DNA | NanR-dimer_1_/DNA | NanR-dimer_2_/DNA | NanR-dimer_2_/DNA | NanR-dimer_3_/DNA | NanR-dimer_3_/DNA |
| * These are the sedimentation and diffusion coefficients observed following 2DSA-Monte Carlo analysis. Parameters are obtained from integration of pseudo-3D plots using UltraScan^15^. All measured values represent the mean from the Monte Carlo analysis. The values in parentheses are the 95% confidence intervals from the Monte Carlo analysis. | | | | | | | | | |
| ^†^ N/D = not determined as species were present within a reaction boundary. | | | | | | | | | |
| ^‡^ Partial concentration is determined from peak integration of the co-migrating species in both the NanR and DNA datasets. Because the data is scaled to molar concentrations, the molar ratio and thus stoichiometry of the hetero-complex can be inferred. | | | | | | | | | |
| ^§^ Partial specific volume ($\bar{v}$) of the hetero-complex, estimated from the weight-average of the protein and DNA components. A $\bar{v}$ of 0.7295 mL g^-1^ was used for NanR, while a $\bar{v}$ of 0.55 mL g^-1^ was used for DNA. The equation used to calculate the weight-averaged $\bar{v}$ is presented in the Methods. | | | | | | | | | |
| ^¶^ The measured molar mass is estimated based on the hydrodynamic parameters (sedimentation and diffusion coefficient) and the weighted-averaged $\bar{v}$, calculated using the measured molar ratio. The values in parentheses are the 95% confidence intervals from the Monte Carlo analysis for the pure components. | | | | | | | | | |
| ^#^ The theoretical mass is predicted based upon the amino acid or nucleic acid sequence within UltraScan^15^. The mass of each hetero-complex is predicted based on the observed molar ratio. | | | | | | | | | |

***Extended Data* Table 6 | X-ray crystallography data collection, refinement and validation statistics***

| **Data collection parameters** | **C-terminal substructure** | **NanR**  (PDB-60N4) |
| --- | --- | --- |
| Beamline | MX2 | MX2 |
| Detector | EIGER x 16M | EIGER x 16M |
| X-ray wavelength (Å) | 1.2781 | 0.9537 |
| Crystal-to-detector distance (mm) | 245 | 200 |
| Space group | *I* 2_1_ 2_1_ 2_1_ | *P* 2_1_ |
| Unit cell *a*, *b*, *c* (Å) | 75.46, 78.35, 88.72 | 39.77, 87.85, 73.87 |
| Unit cell α, β, γ (°) | 90, 90, 90 | 90, 103, 90 |
| Resolution range (Å) | 44.36–2.29 (2.35–2.29) | 43.9–2.10 (2.17–2.10) |
| No. of total reflections | 1,084,309 | 96,673 |
| No. of unique reflections | 22,937 | 28,482 |
| Completeness (%) | 99.6 (97.9) | 98.1 (88.3) |
| *R*_merge_ (%) | 7.7 (49.2) | 4.4 (53.5) |
| CC_anom_ | 44(1) | 0(0) |
| CC_1/2_ (%) | 99.9 (95.2) | 99.9 (69.4) |
| *I*/σ (*I*) | 35.9(5.10) | 15.3 (1.6) |
| **Refinement statistics** |  |  |
| Resolution (Å) |  | 43.9–2.10 |
| No. of reflections used |  | 26,874 |
| No. of reflection used in test set |  | 1,368 |
| *R*_work_/*R*_free_ (%) |  | 18.12/22.95 |
| Total non-H atoms |  | 3,653 |
| Protein/Water |  | 3,586 |
| Zn^2+^ |  | 1 |
| Neu5Ac |  | 66 |
| Mean *B* factor (Å^2^) |  |  |
| Protein |  | 56.5 |
| Zn^2+^ |  | 38.8 |
| Neu5Ac |  | 37.3 |
| Waters/PEG4K |  | 68.1 |
| r.m.s deviations |  |  |
| Bond (Å) |  | 0.009 |
| Angle (°) |  | 1.27 |
| ***MolProbity* statistics†** |  |  |
| Rotamer outliers (%) |  | 3.88 |
| Clashscore |  | 9.17 |
| Ramachandran plot |  |  |
| Favored/allowed regions (%) |  | 98.36/1.64 |
| * Values in parentheses are for the highest resolution shell | | |
| † *MolProbity* structure-validation web service^16^ | | |

***Extended Data* Table 7 | Data collection and analysis statistics from small angle X-ray scattering experiments.**

| **Analysis statistics** | **NanR** | **NanR**  **(+ 20 mM Neu5Ac)** | | | **NanR-dimer_1_/DNA**  **hetero-complex** |
| --- | --- | --- | --- | --- | --- |
| *I*(0) (cm^-1^) (from Guinier analysis) | 0.033 ± 8.9e^-05^ | 0.032 ± 8.9e^-05^ | | | 0.1 ± 5.5e^-04^ |
| *R*_g_ (Å) (from Guinier analysis) | 32.3 ± 0.3 | 31.4 ± 0.2 | | | 33.4 ± 0.6 |
| *I*(0) (cm^-1^) (from *P*(*r*) analysis) | 0.033 ± 0.1e^-03^ | 0.032 ± 0.1e^-03^ | | | 0.010 ± 0.9e^-03^ |
| *R*_g_ (Å) (from *P*(*r*) analysis) | 32.5 ± 0.2 | 31.6 ± 0.2 | | | 32.8 ± 0.2 |
| D_max_ (Å) | 105 | 101 | | | 98 |
| Porod volume (Å^-3^) | 110,038 | 104,710 | | | 108,102 |
| Molar mass (from Porod Volume, kDa) | 64.7 | 61.6 | | | 63.6 |
| Molar mass (from SAXSMoW2*, kDa) | 70.6 | 63.6 | | | 82.9 |
| Calculated dimer MM from sequence (kDa) | 59.0 | 59.0 | | | 70.5 |
| **Data collection parameters** |  | |  | |  |
| Instrument | Australian Synchrotron SAXS/WAXS beamline | | | | |
| Detector | PILATUS 1M (Dectris) | | | | |
| Wavelength (Å) | 1.0332 | | | | |
| Maximum flux at sample | 8 x 10^12^ photons per second at 12 keV | | | | |
| Camera length (mm) | 1600 | | | | |
| *q* range (Å^-1^) | 0.006–0.5 | | | | |
| Exposure time | Continuous 1 second frame measurements | | | | |
| Sample configuration | SEC-SAXS with co-flow | | | | |
| Sample temperature (°C) | 12 | | | | |
| ^a^ http://saxs.ifsc.usp.br/^17^ | | |  |  | |

***Extended Data* Table 8 | Cryo-EM data collection, refinement and validation statistics.**

| **Data collection parameters** | **NanR-dimer_1_/DNA**  **hetero-complex**  (EMD-21652, PDB-6WFQ) | **NanR-dimer_3_/DNA**  **hetero-complex**  (EMD-21661, PDB-6WG7) |
| --- | --- | --- |
| Molecular mass (kDa) | 70.5 | 198.5 |
| Magnification | 215,000 (EFTEM) | 150,000 |
| Voltage (kV) | 300 | 200 |
| Camera | Gatan V2 | FEI Falcon3 |
| Dose rate (e/pixel/s) | 4 | 0.8 |
| Electron exposure (e^-^Å^-2^) | 60 | 45 |
| Defocus range (μm) | -0.5 to -2.5 | -0.5 to -1.5 |
| Pixel size (Å) | 0.68 | 0.94 |
| Symmetry imposed | C1 | C1 |
| Initial particle images (no.) | 695,465 | 957,901 |
| Final particle images (no.) | 141,663 | 48,931 |
| Relative abundance (%)^a^ | 20.39 | 5.11 |
| Applied map sharpening B factor (Å^2^) | -100 | -250 |
| Map resolution (Å) | 3.9 | 8.3 |
| FSC threshold | 0.143 | 0.143 |
| Map resolution range (Å) | 3.710 to 6.096 | N/A |
| **Refinement statistics** | | |
| Model composition |  |  |
| Non-hydrogen atoms | 4164 |  |
| Protein residues | 447 |  |
| Nucleotide residues | 30 |  |
| r.m.s deviations |  |  |
| Bond lengths (Å) | 0.011 |  |
| Bond angles (°) | 1.159 |  |
| **Validation statistics** | | |
| *MolProbity*† score | 2.89 |  |
| Rotamer outliers (%) | 1.09 |  |
| Clashscore, all atoms | 49.60 |  |
| Ramachandran plot |  |  |
| Favored (%) | 83.1 |  |
| Allowed (%) | 16.44 |  |
| Disallowed (%) | 0.46 |  |
| ^a^ Relative abundance with respect to the total number of images in the initial cryo-EM dataset. | | |
| † *MolProbity* is a structure-validation web service^16^ | | |

***Extended Data* References**

1.	Schuck, P. Size-distribution analysis of macromolecules by sedimentation velocity ultracentrifugation and lamm equation modeling. *Biophys. J.* **78**, 1606-19 (2000).

2.	Sievers, F. et al. Fast, scalable generation of high-quality protein multiple sequence alignments using Clustal Omega. *Mol. Syst. Biol.* **7**(2011).

3.	Robert, X. & Gouet, P. Deciphering key features in protein structures with the new ENDscript server. *Nucleic Acids Res.* **42**, W320-W324 (2014).

4.	Little, M.S., Pellock, S.J., Walton, W.G., Tripathy, A. & Redinbo, M.R. Structural basis for the regulation of beta-glucuronidase expression by human gut Enterobacteriaceae. *Proc Natl Acad Sci U S A* (2017).

10.	Lawson, C.L. & Hanson, R.J. *Solving Least Squares Problems*, (Prentice-Hall, Inc., Englewood Cliffs, NJ., 1974).
